# Alien Species of Fish in the Littoral of Volga and Kama Reservoirs (Results of Complex Expeditions of IBIW RAS in 2005-2017)

**DOI:** 10.1101/355362

**Authors:** D.P. Karabanov, D.D. Pavlov, M.I. Bazarov, E.A. Borovikova, Yu.V. Gerasimov, Yu.V. Kodukhova, A.K. Smirnov, I.A. Stolbunov

## Abstract

The paper provides information on alien species of fish caught in the coastal waters of the Volga and Kama river reservoirs. The material was collected during complex ship expeditions of the IBIW RAS in 2005-2017. We have identified habitats and estimated the relative abundance of mass alien species of the Volga-Kama region.

## INTRODUCTION

The problem of the penetration and naturalization of living organisms beyond their historical ranges has not lost its relevance for more than half a century. Human activity plays a significant role in this process. The ever-increasing anthropogenic transformation of the natural environment, combined with global climatic changes, which have intensified since the last decades of the 20th century, have accelerated the transformation of many plant and animal species’ ranges. Man not only conducts mass acclimatization of specific species (cultivation of potatoes, aquaculture of salmonids), but also causes accidental introductions (transfer of invertebrates with ballast water, accidental introductions of fish during the acclimatization of aquaculture objects). The conditions of hydrobionts’ habitat change as a result of human activities, providing a possibility for an increase of some species’ ranges (resulting, for example, in the expansion of the Ponto-Caspian sprat after the construction of reservoirs cascade on Volga).

A quarter of a century ago, in 1992, an international “Convention on Biological Diversity” (CBD), ratified by the Russian Federation in 1995, was signed in Rio de Janeiro. In accordance with paragraph 8 (h) of the CBD, the participating countries are obliged “to prevent introductions, control or destroy those alien species that threaten ecosystems, habitats or species”. In the development of these decisions, the “6th Conference of the Parties to the CBD (Decision VI / 23, 2002, The Hague) approved the” Guidelines for the Prevention of Introduction and Reduction of the Impacts of Alien Species that Threaten Ecosystems, Habitats or Species “. No less attention is paid to the alien species in the “Strategic Plan 2011-2020” of the CBD (Decision X / 2, 2010, Nagoya) and was considered by a separate item at the 12th Conference of the Parties to the CBD (Decision XII / 16, 2014, Pyeongchang). The consistent fulfillment of Russia’s obligations under the CBD is reflected in the successively implemented “National Strategy and Action Plan for the Conservation of Biological Diversity” (Moscow, RF Ministry of Natural Resources, 2002), where the problem of biological invasions is among the main ones. All these approaches are assigned in the “Environmental Doctrine of the Russian Federation”, approved by the RF Government Decree No. 1225-p (2002). As noted in the Fifth National Report “Biodiversity Conservation in the Russian Federation 2014” (Moscow, MNR RF, 2015), the intensity of invasions of alien species of plants and animals in terrestrial and marine ecosystems continues to grow, and one of the main current and promising threats to biological diversity of Russia are invasions of alien species (access to all CBD documents is available at http://www.cbd.int/).

Thus, the determination and timely detection of new species beyond their historical ranges, pathways and vectors of biological invasions is important both theoretically and practically. The creation of a system of reservoirs in the largest European rivers led to a significant transformation of native communities and the formation of conditions favorable for the expansion of the range of different species of hydrobionts. Of course, the study that we conducted can not be considered comprehensive, and the listed species list reflects only the mass alien species that live in the Volga and Kama littoral. At the same time, even according to the presented data, it is possible to trace some regularities in the transformation of local fish communities in the result of alien species introduction, and also to suggest the ways and causes that contributed to the successful expansion of the range of some fish species.

This publication is a continuation and development of work on the inventory of alien species of fish and their role in the ichthyofauna of the Volga reservoirs associated with the scientific activity of Valentina Ivanovna Kiyashko (IBIW RAS), many of which, unfortunately, failed to be realized due to the sudden death of this distinguished researcher.

## MATERIAL AND METHODS

Alien fish species were caught during the annual complex biological expeditions of the IBIW RAS on RV “Akademik Topchiev” in the summer field seasons from 2005 to 2017. The sampling stations were chosen based on the coastal conditions: the presence of shallow water with beds of aquatic vegetation and, if possible, the absence of strong currents. The network of stations was formed with the maximum coverage of the entire water area of each reservoir. The main fishing gear were: beach seine with a size of 10×1.5 m, 4 mm mesh in the cod-end and wings; beach seine with dimensions of 25×1.7 m, 10 mm mesh in wings, 5 mm in the cod-end; square fish lift net with 1.5 m side, 4 mm mesh and ichthyological scoop-net with 4 mm mesh. In total, 276 hauls were performed at 89 stations, not less than 3 passes on each with a distance of 30-50 m and maximum opening of the fishing gear. If possible, the catch was analyzed at the site of fishing. The map of sampling stations is shown in Figure 1. The species of fish and their size and weight characteristics were determined according to the traditional method (Pravdin, 1966). Most of the catch was released back into the reservoir with minimal damage. Thus, more than 19 thousand fish were collected. Alien species were totally fixed in 95% ethanol for further laboratory study. The species were identified using the following keys [Koblitckaya, 1981; Atlas presnovodnykh …, 2003a, b; Kottelat, Freyhof, 2007]. Due to significant changes in the taxonomy of fish, the classification and Latin names here and below are not presented in the original author’s writing, but according to the latest edition of the FishBase database [Froese, Pauly, 2017], macrosystematics according to the latest edition of “Fishes of the World” [Nelson et al., 2016].

**Figure 1.**
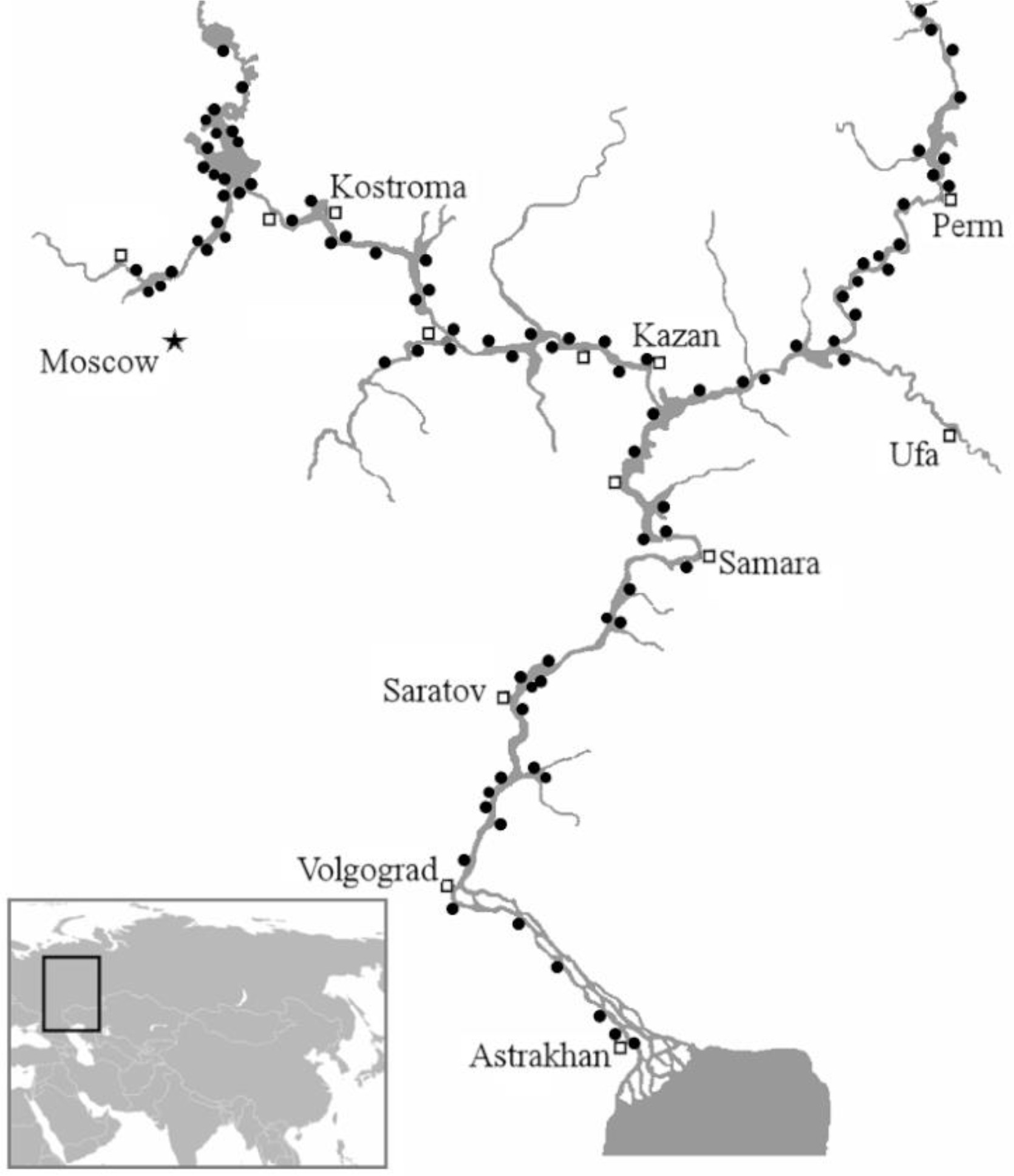
Sampling sites in the Volga-Kama region (2005-2017).

## RESULTS

A total of 124 to 140 fish species inhabit the waterbodies of the studied basin (according to various systematic reports). This variation is mainly conditioned by the continuing process of everlasting review of the taxonomic status of many species and subspecies rather than the actual state of affairs [Mina et al., 2006]. However, even in the case of conservative approach in our littoral catches, the share of alien species in the reservoirs of the Volga ranges from 8% to 32%. For Kama reservoirs, the proportion of alien species is much less - from 2 to 16% (table). Thus, invasive fish are a stable (though often small) component of coastal communities. The fish population of the Volga-Kama region is based on the representatives of the Ponto-Caspian freshwater and boreal plain ichthyofauna. Alien fishes originate mainly from the Ponto-Caspian marine faunistic complex. The most well-represented here – family Gobiidae, from which we found six representatives. It should also be noted that the pelagic area of virtually all water reservoirs is dominated by another Ponto-Caspian “southern invader” – the common kilka [Karabanov, 2013a (Karabanov, 2013a)]. For northern reservoirs, the “northern invader”, representative of the Arctic freshwater faunal complex, the European vendace (Borovikova, 2009), is a permanent component of the pelagic community. Undoubtedly, mass alien fish species have their adaptations that have ensured the success of the introduction, so it is worthwhile to consider the features of the biology of each individual species, separately.

Class: Actinopterygii.

Order: Perciformes.

Family: Gobiidae Cuvier, 1816.

Gobiids is the most well-represented fish family among European invasive fishes. Systematics of gobiids is extremely intricate and validity of some taxa, even when proven using methods of DNA analysis, requires further studying [Nielson, Stepien, 2009; Sorokin et al., 2011]. We found six invasive representatives from this family in the Volga’s basin and only two in the Kama’s basin.

*Ponticola gorlap* (Iljin, 1949) – Caspian bighead goby.

Previously known as *Neogobius iljini* Vasil’eva and Vasil’ev, 1996. According to the latest revision this taxon has been identified as a younger synonym and *P. gorlap* is the accepted valid name [Neilson, Stepien, 2009]. This is the largest of gobiids settling in the Volga’s basin. This species was present in our catches from the Lower Volga up to the Kuybyshev reservoir. The donor region of this fish is probably located in the Lower Volga and its delta as well as in the brackish area of the Caspian Sea. According to literature, Caspian bighead goby penetrated into the Volgograd reservoir in 1970’s (Gavlena, 1977), a decade later – Saratov reservoir (Kozlovskaya, 1997), its expansion in the Kuybyshev reservoir continues [Galanin, 2012]. It is locally present in the Cheboksary reservoir, single specimens were found in the Gorky reservoir (Klevakin at al., 2005). Findings of this species in the Upper Volga reservoirs in 1990’s (Ekologicheskie problemy…, 2001) requires additional verification (Atlas presnovodnykh…, 2003b). There are data [Skomorokhov, 2016] indicating a population of this species in the Moskva River. We did not find it in the Kama reservoirs; however, the probability of its expansion in this direction is very high.

*Ponticola syrman* (Nordmann, 1840) – Syrman goby.

Previously included in the *Neogobius* genus as a separate Caspian subspecies *N. syrman eurystomus* (Atlas presnovodnykh…, 2003b). Extensive phylogenetic study [Neilson, Stepien, 2009] substantiated the isolation of a separate genus *Ponticola* and Syrman goby was considered a representative of this genus. After the return of the systematics of Caspian gobies to dichotomous keys, based on the topology of seismosensory system channels, identified by B.S. Ilin at the beginning of the 20th century and a thorough revision of all Caspian gobies (Opredelitel ryb …, 2013) the modern name *P. syrman* was proposed.

In our catches, Syrman goby is found only on the Lower Volga in the region of Astrakhan. Further studies are required to consider this species as invasive in the Lower Volga. It is known that Syrman goby does not venture upstream of the estuary zones [Freyhof, 2011] and its distribution is limited to the Volga’s delta (Opredelitel ryb., 2013). Finding of Syrman goby in Astrakhan may be explained by their self-dispersion from the Volga’s delta or accidental introduction with ballast waters of marine vessels. It is possible that Syrman goby has formed a local population in Astrakhan area as one of the caught individuals was adult female and another – juvenile underyearling.

*Neogobius melanostomus* (Pallas, 1814) – Round goby.

One of the most widespread invasive species. The origin of the Volga populations can be related both to self-dispersion from the Lower Volga (Gavlena, 1970) and to accidental introductions from the Don delta as a result of acclimatization events (Tsyplakov 1974). At the moment, this species has successfully invaded all reservoirs of Volga [Karabanov et al., 2014]. As for Kama River, round goby was found up to the headwaters of the Votkinsk water reservoir.

The success of the invasion of this alien is due, *inter alia*, to its extraordinary eurybionticity. Round goby can live, reproduce and develop in a wide range of water temperature, oxygen concentration, hydrochemical composition, has a wide feeding spectrum, and is also able to maintain the population size due to early maturation, oogenesis features and high spawning efficiency. At the same time, we should not exclude the essential role of navigation in the distribution of this species. In addition to transporting fish with ballast water and transportation with building materials (sand, gravel), passive transfer of eggs glued to fouling on the bottom of vessels is also possible, which is facilitated by the morphophysiological features of embryogenesis of the round goby (Moskalkova, 1997). All these adaptations determine the possibility of large scale dispersal of round goby in the reservoirs, and the limiting factor is likely to be only the presence of suitable stony-sandy bottoms [Stolbunov et al., 2013].

*Neogobius fluviatilis* (Pallas, 1814) – Monkey goby.

The historical range consists of rivers and brackish areas of the Azov, Caspian and Black seas basins. After dam construction, this species began to expand up the Volga (Caspian populations were likely to be donors). Since the 1970’s was found in the ichthyofauna of Volgograd, in 1980’s – in the Saratov reservoirs [Evlanov et al., 1998]. At the beginning of the 21^st^ century it appeared in the Kuibyshev reservoir, also there are data on single finds in the Cheboksary reservoir [Klevakin et al., 2005 (Klevakin at al., 2005)]. According to our catches, the monkey goby is often found in the reservoirs of the Lower Volga. The northermost catch of a single specimen was made in the upper part of the Kamsky reach of the Kuibyshev water reservoir. This species is probably more widely represented in the reservoirs of the Volga-Kama cascade, but this information requires verification [Shakirova, 2015].

*Proterorhinus sp.* – Tubenose gobies.

The characteristic features of this genus are the following: the anterior nasal cavities are extended into cirriform tubules hanging downward over the upper lip; gill covers are bare, except for their upper part; the bases of the pectoral fins and the back of the throat are covered with cycloid scales [(Atlas presnovodnykh …, 2003b]. The taxonomy of these gobies is also extremely complicated. Different authors indicate the belonging of the Volga tubenose gobies to the species *Pr.* cf. *semipellucidus* [Neilson, Stepien, 2009], *Pr. nasalis* [Sorokin et al., 2011], *Pr. marmoratus* sensu lato [Galanin, 2012], *Pr. semilunaris* [Slynko et al., 2013]. The issue of the donor region and the taxonomic belonging of tubenose gobies of the Volga-Kama basin, undoubtedly, requires a separate study [ (Opredelitel ryb …, 2013].

Stump-necked gobies, as well as the gobies-rounds are the most common invaders of the Volga. This species was first discovered in the 1980’s in the Saratov Reservoir [Evlanov et al, 1998 (Evlanov et al., 1998)]. Later, probably because of their small size, these gobies have quietly settled in all reservoirs of the Volga, and also creating a population in the Moscow River [Sokolov, Tsepkin, 2000); Klevakin at al., 2005; Ryby Rybinskogo …, 2015]. According to our catches, the tubenose goby is present in virtually all Volga’s reservoirs, reaching a considerable abundance, in some cases (table).

*Benthophilus stellatus* (Sauvage, 1874) – Stellate tadpole goby.

Stellate tadpole goby is one of the most poorly studied species, the natural range of which covers brackish limans, gulfs, rivers and coastal lakes of the Black, Azov and Caspian seas basins [Atlas presnovodnykh …, 2003b]. DNA-barcoding of these fish [Kodukhova et al., 2016] did not confirm the idea that Volga’s population of stellate tadpole goby is represented by Don River species *B. durrelli* [Boldyrev, Bogutskaya, 2007]. By now, stellate tadpole goby is regularly found in the reservoirs of the Lower and Middle Volga [Kasyanov, Klevakin, 2011; Shakirova et al., 2015], in the Cheboksary reservoir within the territory of Nizhniy Novgorod area (Klevakin et al., 2005), and a single adult specimen was caught in the Rybinsk Reservoir [Kodukhova et al., 2016].

The settlement of the stellate tadpole goby in the reservoirs of the Middle Volga, apparently, is associated with large-scale construction of dams and hydroconstructions in the second half of the twentieth century. Undoubtedly, the population from the Kuibyshev reservoir serves as the donor region for the resettlement of the star-shaped head for the reservoirs of the Upper Volga. The slight genetic differentiation of the Middle and Upper Volga specimens from the Black Sea fish probably indicates the origin of the populations of the stellate tadpole goby of the Kuibyshev Reservoir as a result of accidental introduction of fish together with mysids from the Azov-Black Sea basin [Gavlena 1973] (Figure 2.c). The genetic variant from the Saratov reservoir differs from all other sequences and, probably, has a different origin: taking into account that this reservoir is the closest of all the above to the Caspian Sea, the population of the stellate tadpole goby that resides there can have be of Caspian origin [Kodukhova et al., 2016]. Undoubtedly, further phylogeographic studies of the freshwater populations of these fish are required.

**Figure 2.**
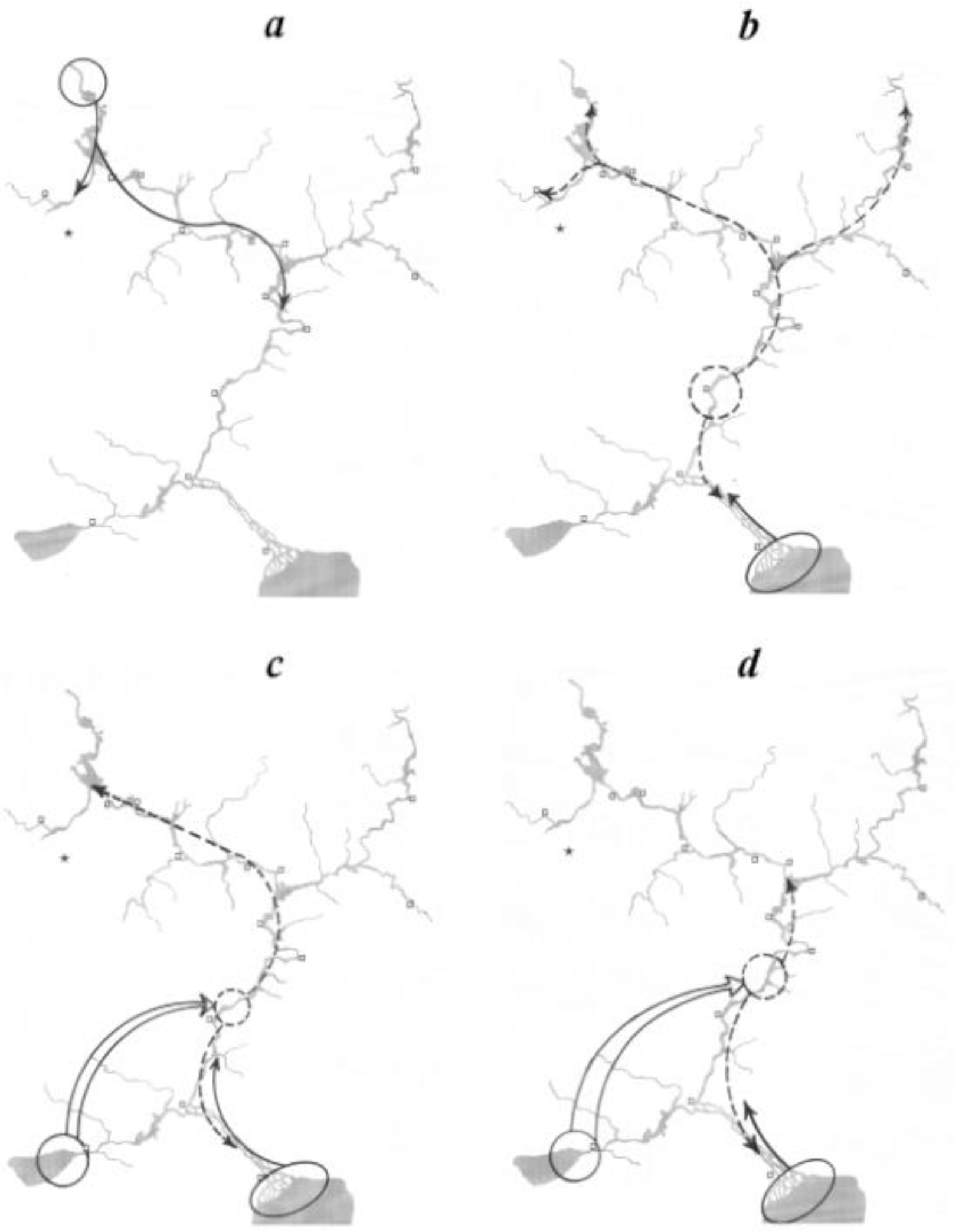
Probable donor regions, pathways of penetration and range expansion of mass invasive fish species in the Volga-Kama region. ***a*** – vendace, *Coregonus albula*; ***b*** - common kilka, *Clupeonella cultriventris*; ***c*** - stellate tadpole goby, *Benthophilus stellatus*; ***d*** – black-striped pipefish, *Syngnathus abaster.*

**Table 1.**
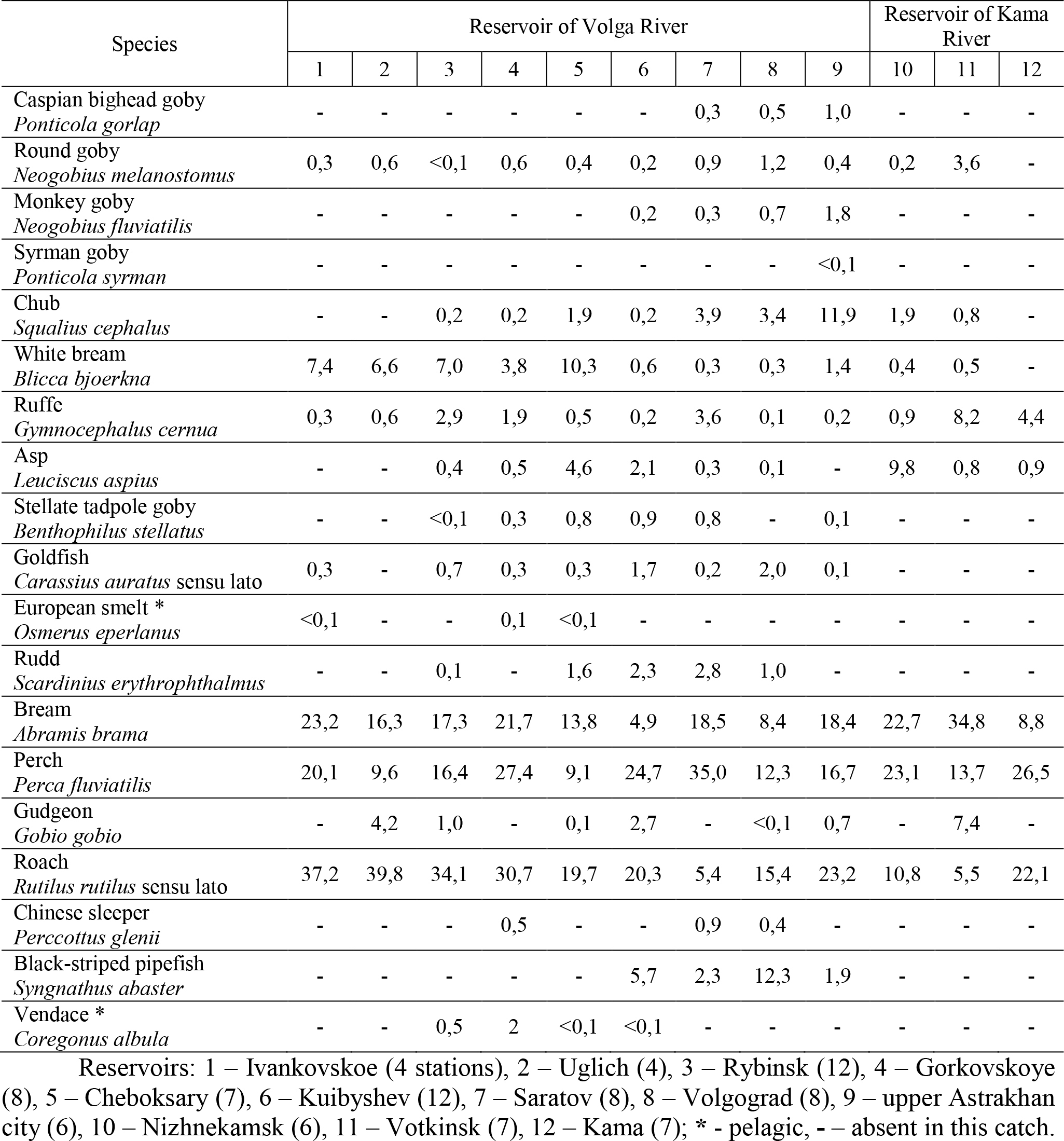
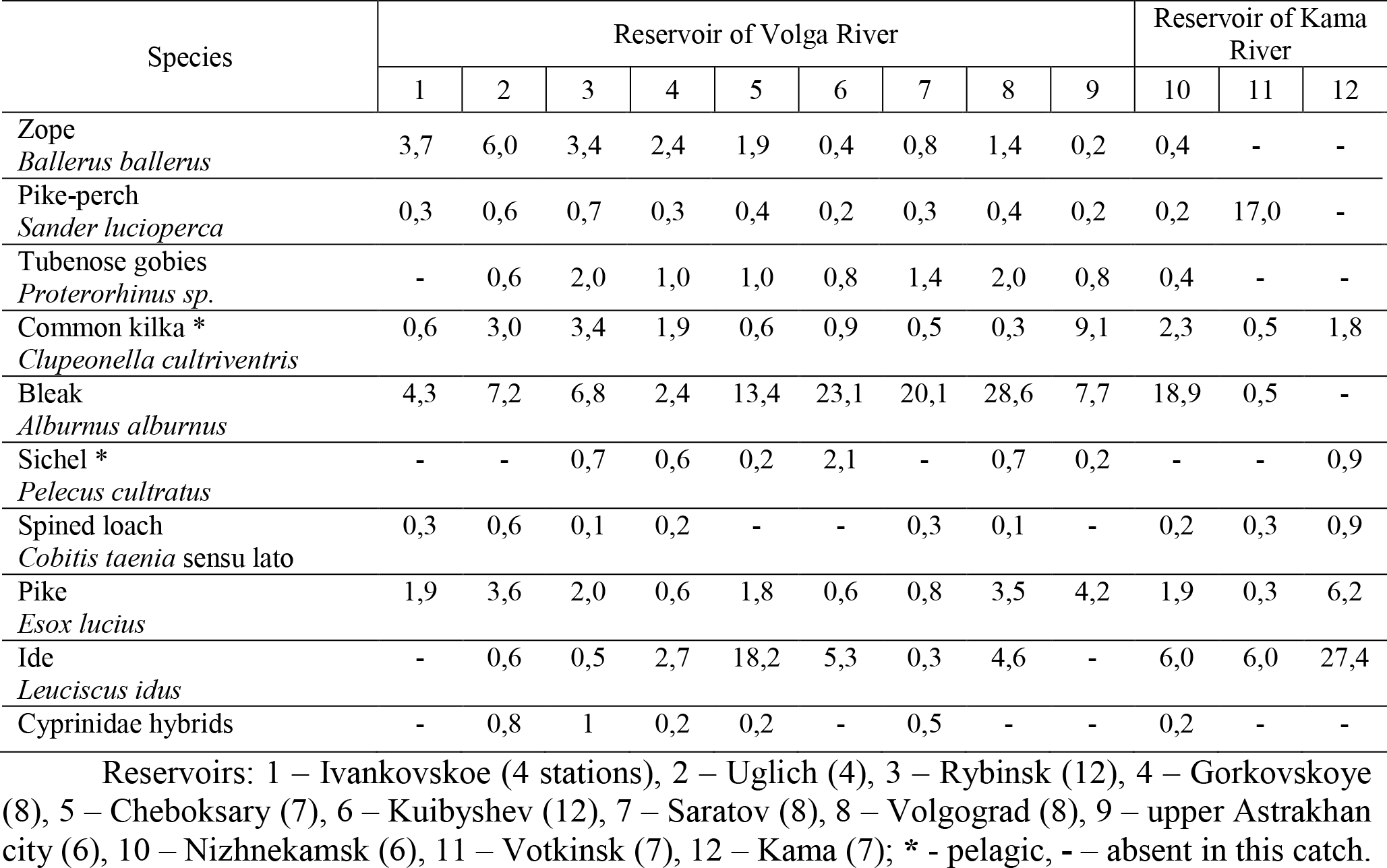
The ratio of various mass invasive fish species in 2005-2017 catches.

| Species | Reservoir of Volga River |  |  |  |  |  |  |  |  | Reservoir of Kama River |  |  |
| --- | --- | --- | --- | --- | --- | --- | --- | --- | --- | --- | --- | --- |
|  | 1 | 2 | 3 | 4 | 5 | 6 | 7 | 8 | 9 | 10 | 11 | 12 |
| Caspian bighead goby<br>*Ponticola gorlap* | - | - | - | - | - | - | 0,3 | 0,5 | 1,0 | - | - | - |
| Round goby<br>*Neogobius melanostomus* | 0,3 | 0,6 | <0,1 | 0,6 | 0,4 | 0,2 | 0,9 | 1,2 | 0,4 | 0,2 | 3,6 | - |
| Monkey goby<br>*Neogobius fluviatilis* | - | - | - | - | - | 0,2 | 0,3 | 0,7 | 1,8 | - | - | - |
| Syrman goby<br>*Ponticola syrman* | - | - | - | - | - | - | - | - | <0,1 | - | - | - |
| Chub<br>*Squalius cephalus* | - | - | 0,2 | 0,2 | 1,9 | 0,2 | 3,9 | 3,4 | 11,9 | 1,9 | 0,8 | - |
| White bream<br>*Blicca bjoerkna* | 7,4 | 6,6 | 7,0 | 3,8 | 10,3 | 0,6 | 0,3 | 0,3 | 1,4 | 0,4 | 0,5 | - |
| Ruffe<br>*Gymnocephalus cernua* | 0,3 | 0,6 | 2,9 | 1,9 | 0,5 | 0,2 | 3,6 | 0,1 | 0,2 | 0,9 | 8,2 | 4,4 |
| Asp<br>*Leuciscus aspius* | - | - | 0,4 | 0,5 | 4,6 | 2,1 | 0,3 | 0,1 | - | 9,8 | 0,8 | 0,9 |
| Stellate tadpole goby<br>*Benthophilus stellatus* | - | - | <0,1 | 0,3 | 0,8 | 0,9 | 0,8 | - | 0,1 | - | - | - |
| Goldfish<br>*Carassius auratus sensu lato* | 0,3 | - | 0,7 | 0,3 | 0,3 | 1,7 | 0,2 | 2,0 | 0,1 | - | - | - |
| European smelt *<br>*Osmerus eperlanus* | <0,1 | - | - | 0,1 | <0,1 | - | - | - | - | - | - | - |
| Rudd<br>*Scardinius erythrophthalmus* | - | - | 0,1 | - | 1,6 | 2,3 | 2,8 | 1,0 | - | - | - | - |
| Bream<br>*Abramis brama* | 23,2 | 16,3 | 17,3 | 21,7 | 13,8 | 4,9 | 18,5 | 8,4 | 18,4 | 22,7 | 34,8 | 8,8 |
| Perch<br>*Perca fluviatilis* | 20,1 | 9,6 | 16,4 | 27,4 | 9,1 | 24,7 | 35,0 | 12,3 | 16,7 | 23,1 | 13,7 | 26,5 |
| Gudgeon<br>*Gobio gobio* | - | 4,2 | 1,0 | - | 0,1 | 2,7 | - | <0,1 | 0,7 | - | 7,4 | - |
| Roach<br>*Rutilus rutilus sensu lato* | 37,2 | 39,8 | 34,1 | 30,7 | 19,7 | 20,3 | 5,4 | 15,4 | 23,2 | 10,8 | 5,5 | 22,1 |
| Chinese sleeper<br>*Perccottus glenii* | - | - | - | 0,5 | - | - | 0,9 | 0,4 | - | - | - | - |
| Black-striped pipefish<br>*Syngnathus abaster* | - | - | - | - | - | 5,7 | 2,3 | 12,3 | 1,9 | - | - | - |
| Vendace *<br>*Coregonus albula* | - | - | 0,5 | 2 | <0,1 | <0,1 | - | - | - | - | - | - |
|  | 1 | 2 | 3 | 4 | 5 | 6 | 7 | 8 | 9 | 10 | 11 | 12 |
| Zope<br>*Ballerus ballerus* | 3,7 | 6,0 | 3,4 | 2,4 | 1,9 | 0,4 | 0,8 | 1,4 | 0,2 | 0,4 | - | - |
| Pike-perch<br>*Sander lucioperca* | 0,3 | 0,6 | 0,7 | 0,3 | 0,4 | 0,2 | 0,3 | 0,4 | 0,2 | 0,2 | 17,0 | - |
| Tubenose gobies<br>*Proterorhinus sp.* | - | 0,6 | 2,0 | 1,0 | 1,0 | 0,8 | 1,4 | 2,0 | 0,8 | 0,4 | - | - |
| Common kilka *<br>*Clupeonella cultriventris* | 0,6 | 3,0 | 3,4 | 1,9 | 0,6 | 0,9 | 0,5 | 0,3 | 9,1 | 2,3 | 0,5 | 1,8 |
| Bleak<br>*Alburnus alburnus* | 4,3 | 7,2 | 6,8 | 2,4 | 13,4 | 23,1 | 20,1 | 28,6 | 7,7 | 18,9 | 0,5 | - |
| Sichel *<br>*Pelecus cultratus* | - | - | 0,7 | 0,6 | 0,2 | 2,1 | - | 0,7 | 0,2 | - | - | 0,9 |
| Spined loach<br>*Cobitis taenia sensu lato* | 0,3 | 0,6 | 0,1 | 0,2 | - | - | 0,3 | 0,1 | - | 0,2 | 0,3 | 0,9 |
| Pike<br>*Esox lucius* | 1,9 | 3,6 | 2,0 | 0,6 | 1,8 | 0,6 | 0,8 | 3,5 | 4,2 | 1,9 | 0,3 | 6,2 |
| Ide<br>*Leuciscus idus* | - | 0,6 | 0,5 | 2,7 | 18,2 | 5,3 | 0,3 | 4,6 | - | 6,0 | 6,0 | 27,4 |
| Cyprinidae hybrids | - | 0,8 | 1 | 0,2 | 0,2 | - | 0,5 | - | - | 0,2 | - | - |

Reservoirs: 1 – Ivankovskoe (4 stations), 2 – Uglich (4), 3 – Rybinsk (12), 4 – Gorkovskoye (8), 5 – Cheboksary (7), 6 – Kuibyshev (12), 7 – Saratov (8), 8 – Volgograd (8), 9 – upper Astrakhan city (6), 10 – Nizhnekamsk (6), 11 – Votkinsk (7), 12 – Kama (7); \* - pelagic, - – absent in this catch.

Family: Odontobutidae Hoese et Gill, 1993 – Freshwater sleepers.

In the basin of the Volga and Kama (but not in these rivers themselves) there is only one species - a rotan-“headdress”.

*Perccottus glenii* Dybowski, 1877 – Chinese (Amur) sleeper.

Amur sleeper is one of the most mass alien fish species in Europe [Reshetnikov, Ficetola, 2014]. Previously belonged to the family Eleotrididae, but assigned into a separate taxon during the last revision [Nelson et al., 2016]. Inhabits small lakes, ponds, oxbows. Almost absent in large waterbodies. In extremely rare cases, this species is observed in shallow, overgrown areas of Volga River reservoirs [Kasyanov, Goroshkova, 2012]. There are data [Shakirova et al., 2015] on the existence of a large number of Amur sleepers populations in the Staromaynsky Bay of the Kuibyshev Reservoir. Also, according to the literature [Semenov, 2010 (Semyonov, 2010)], the Amur sleeper is sporadically found in many overgrown bays and inundations of the Kuibyshev reservoir, but not in the open water area and the course. To date, we found only three biotopes in the Volga, where Amur sleeper is always encountered. The first biotope is an overgrown shallow gulf across from Barminsky island (Cheboksary water reservoir within the Nizhny Novgorod region, 56 ° 15 ’N, 45 ° 50’ E) in a channel to the left of the fairway. Six fish with a body length of 33-108 mm were caught in this area. A large number of young cyprinid fish (bleak, roach, bream) as well as juvenile perch were caught here together with Amur sleeper. It is possible that Amur sleeper enters the given bay through a system of channels and ditches from the adjacent floodplain ponds. The second discovered biotope is a shallow, heavily overgrown bay of the discharge channel of the Balakovo NPP (Saratov Reservoir, 42°12’N, 47°94’E). Here we caught three 0+ and one 1+ fish. The depth at this site was less than 1 m the current is practically absent. This site was also inhabited by 0+ perch and bleak. The third biotope is the bay of river Kurdyum (Volgograd reservoir, upstream of Saratov city, 51°40′N, 46°10′E). Here we caught five Amur sleepers with a body length of 45-55 mm. Apart from the Amur sleeper the catch also included cyprinid and perch fry and also invasive fish: tubenose goby and black-striped pipefish.

Thus, both within the native and acquired range, Amur sleeper does not actually occur in large reservoirs and main courses of rivers. It is obvious that in these waters there are certain environmental factors that prevent the existence of this species. Most often the high vulnerability of Amur sleeper to predators, as well as the complete avoidance of waterbodies with even average flow velocities are indicated among such factors [(Ryby v zapovednikakh …, 2010].

Order Salmoniformes

Family: Salmonidae Rafinesque, 1815.

One representative of salmonids – European vendace demonstrates the biggest acquired range. Despite multiple attempts of acclimatizing other salmonid fishes, none of them yielded significant results (Ryby Rybinskogo…, 2015).

*Coregonus albula* (Linnaeus, 1758) – European vendace.

Earlier, European vendace (as well as other coregonids) belonged to family Coregonidae. According to the latest revision, coregonids are now considered a separate subfamily Coregoninae and belong to the family Salmonidae [Nelson et al., 2016]. Traditionally, the vendace is considered as an example of a “northern” invader, which became a part of Volga’s ichthyofauna due to migration from the Lake Beloye (Figure 2.a), although the intended introduction of the vendace (and other coregonids) into the reservoirs of the Upper Volga from other regions is not to be ruled out [Ryby Rybinskogo …, 2015]. Identification of the taxonomic status and modes of formation of the population structure of European vendace in the European Russia, as well as the nature of the currently significant level of its genetic variability, requires special study [Borovikova et al., 2013]. A separate issue is the effectiveness of acclimatization measures of coregonids and their impact on the gene pool of native and new populations. It is known that the representatives of not only different species, but even the genera of this subfamily, easily interbreed with each other, giving fertile offspring [(Borovikova, Makhrov, 2013]. In conditions when numerous forms and species were introduced into the Volga, hybridization (including introgressive) could have occurred rather frequently, although in the case of coregonids, as well as other taxa, the total absence of acclimatization measures’ effects cannot be ruled out. Vendace is a very rare catch in the shallows of Volga reservoirs, although in the open area it is a small but stable component of the pelagic part of the fish community. The southernmost point where we caught vendace is the upper reach of Zhigulyovskaya hydroelectric power station dam near the city of Togliatti.

Order: Osmeriformes.

Family: Osmeridae.

The only representative of this family – European smelt inhabits the Upper Volga basin.

*Osmerus eperlanus* (Linnaeus, 1758) – European smelt.

European smelt is an ordinary inhabitant of reservoirs of the North-West of Russia. There is a large (smelt) and small (snetok) form [Atlas presnovodnykh …, 2003a]. Undoubtedly, smelt (snetok) penetrated into the Volga water reservoirs from Lake Beloye and in the second half of the 20th century has formed a large population in the Rybinsk Reservoir with significant interannual fluctuations in abundance, which is also characteristic of other short-cycle species [Ryby Rybinskogo …, 2015]. Migrating downstream of Volga, the smelt settled up to the Kuibyshev reservoir, where it also created a population with extremely high interannual variations in population (Tereshchenko, Tereshchenko, 2017). Currently, smelt is only rarely found in some reservoirs of the Upper Volga, while the main factor in the distribution of smelt is the thermal dynamics of the water masses of the reservoir [Ryby Rybinskogo …, 2015]. Thus, in the Rybinsk Reservoir, its catches are confined mainly to Sheksna reach, and several of the 0+ fish were caught in the upper reaches of the Ivankovo Reservoir in the cold summer of 2017.

OTpяД: Syngnathiformes.

CeMeйcTBo: Syngnathidae Rafinesque, 1810.

The only representative of this family to inhabit Volga’s basin is the black-striped pipefish, especially widespread in the Lower Volga.

*Syngnathus abaster* Risso, 1827 – black-striped pipefish.

Earlier pipefish found in Volga belonged to *S. nigrolineatus* species. At present, this taxon is considered to be the junior synonym of *S. abaster* [Eschmeyer et al., 2017], although the definition of taxonomic status for large population groups requires a separate study. Based on the study of the genetic variability of pipefish populations in the Caspian and Black seas, the existence of genetically isolated groups has been established, while the Volga pipefish are genetically close to the population of the Black Sea basin probably originating from them[Kiryukhina, 2013]. Thus, apparently, in this case (as well as with stellate tadpole goby), introduction of fish from the Azov-Black Sea basin took place, with the only difference in the vector of pipefish invasion being directed mainly southwards, downstream of Volga (Figure 2 d), while for stellate tadpole goby – to the north. Probably, the peculiarities of pipefish biology, primarily the temperature requirements for spawning and feeding with microplankton, determine the boundaries of the dispersal of this species: for the whole history of observations, only two pipefish were caught in the Cheboksary reservoir (Klevakin et al., 2005 (Klevakin at al., 2005)], whereas in the Lower Volga it is a common and, often, a numerous species in the coastal part of the fish community.

OTpяД: Clupeiformes.

CeMeйcTBo: Clupeidae.

Common kilka, the only clupeid species to inhabit the entire cascade of Volga’s reservoirs, demonstrates the highest abundance and the largest area of acquired range among freshwater clupeids.

*Clupeonella cultriventris* (Nordmann, 1840) – Ponto-Caspian sprat.

Like many other fish species, which significantly expanded their range in a short time, common kilka has also “suffered” from taxonomic perturbations. In the latest revision of clupeids Svetovidov [Svetovidov, 1973] substantiates the suitable nomenclatural name of the species using the older synonym, *Clupeonella cultriventris*, while the status of subspecies (Black Sea-Azov and Caspian kilkas) was put under question [Atlas presnovodnykh …, 2003a]. However, Kottelat in his work utilizes minor morphological differences (Kottelat, Freyhof, 2007) arguing that there are four distinct species of kilkas: the Caspian (*C. caspia*), the Black Sea (*C. cultriventris*), Abrau (*C. abrau*) and freshwater (*C. tcharchalensis*). According to the list of species [Eschmeyer et al., 2017], Volga populations are represented by the latter – *C. tcharchalensis*. At the same time, study of the genetic diversity of common kilka from the whole range [Slynko et al., 2010; Karabanov, 2013a] showed that the Black Sea-Caspian kilka is genetically represented by a single unity in all its modern range, and the allocation of independent taxa does not have a sufficient basis. A certain genetic originality of the freshwater Volga-Kama populations of the common kilka, presumably, is related to the features of their origin from the freshwater resident form from the “Saratov pools” that emerged after the Khvalyn transgression of the Caspian Sea, 40-20 thousand years ago. Thus, the peculiarities of the population-genetic structure of the common kilka are probably a consequence of its origin (which is characteristic of other animals as well), and not of the polyphyletic nature of this taxon [Karabanov, 2013b]. For the final clarification of the status of freshwater kilkas, we performed a DNA-barcoding (the 5 ’region of the mtDNA COI locus), which is the *de facto* standard for the species identification of fish [Ward et al., 2009]. Two unique haplotypes (NCBI GenBank KR075819 and KR075820) were identified in the Upper Volga populations of common kilka (undoubtedly, being representatives of freshwater kilkas). According to the translated amino acid sequence, these haplotypes, the Black Sea haplotype KJ552938, as well as the sequence AP009615 of the total mtDNA and the reference sequence NC_015109 do not differ, and all nucleotide substitutions are synonymous, which may indicate a genetic unity between different populations of common kilka, so that isolation of single species of freshwater kilkas is also not confirmed by DNA-barcoding.

As was shown earlier [(Karabanov, 2013b], the population of the Lower Volga is characterized by a greater balance of the genetic structure in terms of the observed and expected frequencies of allozymes of 17 genetic loci (in the Vologograd reservoir there none, and in Saratov reservoir only 1 locus is characterized by the violation of equilibrium frequencies). When moving up the cascade of reservoirs, the share of loci with a violation of genetic equilibrium gradually increases: from one third for Gorky, to half the loci in the Ivankovo reservoir. Undoubtedly, the key factor in the violation of genetic balance here is the selection, namely, the differentiated survival of various genotypes among the 0+ kilkas during the first winter [Karabanov, 2013a].

At present, common is a dominant species (and often a super-dominant) in the pelagic part of the fish community of Volga reservoirs. This species is characterized by multi-year cycles (6-8 years) of abundance in the reservoirs of the Upper Volga [Kiyashko et al., 2012]. For the reservoirs of the Middle Volga, such powerful “waves of life” are not noted, which is confirmed by both our and literary data. In these waterbodies, common kilka has settled long time ago, has occupied its ecological niche in the pelagic community resulting in the absence of significant demographic fluctuations of these populations.

The extreme point of distribution of self-sustaining populations of common kilka northwards in the Volga-Kama region is the Sheksna Reservoir (Figure 2.b). Only in the summer of 2017, three mature females of this species were caught in Lake Beloye. However, the absence of yearlings and any significant catches of this species does not allow concluding the successful settlement of kilka in this waterbody. Probably, a key role in limiting the distribution of common kilka to the north is played by the change in abiotic (temperature, mineralization, hydrological regime) and biotic factors (quantity and quality of food consumed, availability of food rivals and predators) [Kiyashko et al., 2012]. Another hypothesis [Martemyanov, Borisovskaya, 2010] states that decrease of mineralization in the northern waterbodies along with a decrease in the concentration of sodium in water leads to an increase in the tensions of intracellular systems providing sodium homeostasis and in the degree of kilka’s vulnerability to abrupt changes in the mineralization. This circumstance probably prevents common kilka from spreading northwards (up to Lake Beloye) into freshwater waterbodies with a lower sodium content in the water. The features of hydrology and morphology of Lake Beloye may provide another possible obstacle to the distribution of kilka further north. An assumption of Yu.S. Reshetnikov (oral communication) kilka’s pelagic eggs in Lake Beloye, may become gradually submerged in the bottom layer. Due to the hydrological features in this zone, constant shaking and mixing of the sediments takes place, which leads to mechanical damage and death of the eggs. Concerning biotic factors, one can note the decrease in the number and size of food objects in the Sheksna reservoir in comparison with the Rybinsk and Gorky reservoirs [Ekologicheskie problemy …, 2001]. Also in the more northern reservoirs, the number of food competitors (smelt, vendace and juvenile perch) and the number of predators – zander and large perch is higher, which also complicates the existence of this invasive species.

Mainly aquaculture objects inadvertently fleeing from fish farms are among other alien fish species sporadically encountered in catches in the Volga-Kama region. Among these, first of all, one can note adult individuals of the river eel *Anguilla anguilla* (Linnaeus, 1758), whose larvae were mass released into Lake Seliger from where they could penetrate into Volga [Atlas presnovodnykh …, 2003a]. The same source of new findings of alien fish species explains the capture of the channel catfish *Ictalurus punctatus* (Rafinesque, 1818), brown trout *Salmo trutta* Linnaeus, 1758 and rainbow trout *Parasalmo mykiss* (Walbaum, 1792), etc. Aquaculture of the Far Eastern fish species results in captures of silver *Hypophthalmichthys molitrix* (Valenciennes, 1844) and bighead *Hyp. nobilis* (Richardson, 1845) carps and grass carp *Ctenopharyngodon idella* (Valenciennes, 1844). Unfortunately, many aquaculture farms lack an effective system for the registration and control of aquaculture facilities, which leads to sporadic escape of alien species, and similar facilities become sources of “chronic pollution” by invasive species.

There are data on the disparate populations of the Ukrainian *Pungitius platygaster* (Kessler, 1859) and the nine-spined *Pung. pungitius* (Linnaeus, 1758) sticklebacks in Volga reservoirs [Atlas presnovodnykh …, 2003b]. The latter is also noted for the Kama basin [Askeyev et al., 2010]. However, the current status of populations and the ways of penetration of sticklebacks into these reservoirs require careful verification and a separate study. Finally, another way of increasing the biodiversity of reservoirs is the activity of aquarium enthusiasts. An example of such an exotic invader is an aquarium cyprinodontid guppy fish -*Poecilia reticulata* Peters, 1859. Small populations of this species live only in heated waste and sewage waters of large cities and industrial enterprises. It is possible that the study of such year-round heated areas can lead to the discovery of other tropical aquarium fish [Zworykin, Pashkov, 2010].

## DISCUSSION

There are no doubts that the study and prediction of biological invasions in the waterbodies of Volga-Kama region requires constant and detailed monitoring of the entire water area. In our study, it is impossible to cover such an array of data, and such a task has not been set. Special studies on invasive fish species have been conducted in almost every large waterbody. We also did not consider the issue of mixing species [Levin et al., 2017], or paleoinvasions or phylogenetic lines in roach. A separate topic is the determination of the role of hybrids (primarily cyprinids) as an “alien” component of aquatic communities [Kodukhova, 2011], including those that are important in indicating the quality of the environment. The problem of paleoinvasions and expansion of ranges of already assimilated fish deserves special attention that goes beyond the scope of this study. Such an example can be the expansion of the range of vendace in the south direction from the preglacial refugia located in the European North [Borovikova et al., 2013]. Interestingly, similar processes of settlement from north to south are demonstrated by aquatic invertebrates [Kotov et al., 2016], and the role of such refugia and the northern corridors of self-dispersal of hydrobionts in the formation of biodiversity are strongly underestimated. An example of the expansion of the “local” species is the active development of the *Carassius auratus* sensu lato (whose taxonomic status is still being discussed [Kottelat, Freyhof, 2007, Eschmeyer et al., 2017]) in Volga reservoirs. This process probably takes place due to overall climate warming, an increase in the degree of overgrowth of the littoral and a decrease in predator pressure [Gerasimov et al., 2018].

At the same time, even the available material makes it possible to identify a number of problems associated with the expansion of ranges of hydrobionts beyond their historical limits. The first and most important problem that arises for any researcher is the correct identification and cataloguing of discovered alien species. Even our small sampling volume shows that reliable identification of the invader is complicated due to not only the objective complexity of identification (for example, many diagnostic features in gobiids have overlapping values), but also due to taxonomic innovations caused by the never-ending period of fragmentation in fish systematics [Mina et al., 2006].

The procedure of DNA-barcoding which has now become routine is utilized to solve the problem of species identification [Ward et al., 2009], making it possible to reliably identify fry and very small fish, as well as damaged specimens and those that are in poor condition. Despite the fact that in the Preamble of the “International Code of Zoological Nomenclature” [ICZN, 1999] freedom of thought and actions in the field of taxonomy is declared, constant revisions of the taxonomic status (especially in new parts of the range of invasive fish) create significant difficulties for both the compilers of faunistic lists, and their users. In any case, even the involvement of the DNA-barcoding procedure can not serve as an ultima ratio for revising the taxonomic status of certain groups. First of all, this is due to the very procedure of restoration of the phylogenetic tree. In the case of using one (or even several) genetic loci, the restored “tree of genes” may not reflect the real “species tree”, which can have not only purely mathematical but also biological reasons [Nei, Kumar, 2000; Swenson, El-Mabrouk, 2012; Hellmuth, 2017]. The use of the methods of genosystematics in some cases makes it possible to identify a significant number of cryptic species, for example, both from poorly studied and extremely diverse coral fish [Hubert et al., 2012], and among seemingly well-morphologically studied aquatic invertebrates [Bekker et al ., 2016]. On the other hand, there are hydrobionts, for example, the sea bass *Sebastes* from the North Atlantic well-differing both in terms of morphology and ecology but demonstrating extremely low genetic differentiation [Artamonova et al., 2013], whereas *Daphnia magna* crustacean, possesses extremely high intraspecific genetic diversity [Bekker et al., 2018], often exceeding even the intergeneric limits in other animals. A study of the genetic diversity of fish in North America also revealed the limitations of successful species identification using DNA analysis [April et al., 2011]. Perhaps this problem will be solved with the massive introduction of high-performance sequencing (NGS) technologies and multi-gene sample identification, as well as non-invasive methods for the analysis of “environmental DNA” (eDNA).

At present, it seems that the optimal option is genetic testing of species with further development of dichotomous identification keys for widespread use even by non-specialists. This approach is partially applied in [Opredelitel ryb …, 2013] in the section on Gobiidae. Here the main systematics is based on genetic studies, but the construction of the identification keys is based on classical morphological features.

Another interesting issue related to the settlement of species beyond the limits of historical ranges is the definition of donor regions, the ways of penetration and the direction of the expansion of ranges. Many special studies of both national and foreign research groups are devoted to these problems [reviews: (Biologicheskie invazii …, 2004); (Slynko, Tereshchenko, 2014); Biological invasions …, 2015]. In general, large, regulated rivers are convenient corridors for the dispersal of hydrobionts. The Kuibyshev Reservoir occupies a special place in the Volga cascade. Similar to the Rybinsk reservoir, the Kuibyshev reservoir is characterized by a large area, a diverse morphology of the coast (a multitude of tributaries and large tracts formed by the confluence of large rivers), various biotopes (extensive pelagic,multiple shallows, overgrown and sandy-pebbled littoral) – all this provides a lot of various niches and creates extremely favorable conditions for the acclimatization of new species. Pathways of dispersal of species with latitudinal expansion of ranges along Volga as well as those penetrating into Kama pass through this reservoir. As a result, of these processes, up to a third of the ichthyofauna of the Kuibyshev water reservoir is represented by alien species [Shakirova et al., 2014]. Thanks to this strategic position of the Kuibyshev reservoir, there is a possibility of “mixing” of different phylogenetic lineages of some species of invasive fish (for example, different lineages of pipefish and gobies), which creates additional difficulties for studying their population-genetic structure.

The factors limiting the dispersal of alien species are, first of all, the temperature and the physicochemical parameters of the reservoir. An example of this is the limitation of northbound expansion of pipefish and common kilka and southbound expansion of vendace range. Such relationship is not observed for another invader – the Amur sleeper, which has a wide range of thermoresistance [Golovanov et al., 2013], making it possible for this species to successfully occupy all possible small waterbodies of Europe [Reshetnikov, Ficetola, 2014], and the expansion of its range, is almost entirely limited by biological factors (the presence of predators and natural competitors).

The donor area of many alien fish species of the Volga-Kama region is probably the Azov-Black Sea basin [Biologicheskie invazii …, 2004]. These include, at least for the Middle and Upper Volga, stellate tadpole goby, tubenose goby and pipefish. These species might have been introduced to Volga during massive work on the introduction of forage invertebrates from the Don and the Sea of Azov, or during the river transportation of a large number of building materials and associated cargo in the 1960’s. A number of other self-dispersing species, especially in the Lower Volga (some populations of round goby, Syrman goby and pipefish) originate from estuarine populations of the Volga delta and brackish sections of the Caspian Sea. Common kilka settlement represents a separate mode of invasive processes in the Volga-Kama region. As shown by studies on the genetic structure [Karabanov, 2013a (Karabanov, 2013a)], the origin of the Volga’s kilka is probably associated with the resident form from the “Saratov pools”. Prior to the regulation of the Volga, this little-known freshwater form inhabited the pools and oxbows near the city of Saratov [Svet-ovidov, 1952]. After the creation of reservoir cascade this freshwater kilka was able to settle in the Saratov reservoir, and later along the entire Volga. If we accept the assumption on the origin of the Volga populations from the kilka’s form adapted to the conditions of fresh water, then the high speed of its dispersal becomes understandable. Common kilka in its whole range represents one species both genetically and morphologically. Therefore, the resident kilka of Saratov pools probably represented a freshwater physiological race of *C. cultriventris*. These features, apparently served for (unreasonable, in our opinion), isolation of freshwater kilkas into a separate taxon.

## CONCLUSION

Thus, observing the ongoing processes of species dispersal outside the historical range, this process is studied in three stages: first, the change in the distribution boundaries (independent, associated with human activities or in combination of these causes); secondly, acclimatization and settlement in new habitats; and, thirdly, the completion of the introduction and determination of the invaders niche in the structure of local communities. Changes in the gene pool of invaders may serve as markers of these stages. These processes have been studied most fully for the common kilka [Karabanov, 2013a]. It is known, that both the rapid change in the genotype and the emergence of adaptations without structural rearrangements of the genome is possible for invaders [Prentis et al., 2008]. The available data on genetic processes in newly formed populations of alien species of the Volga-Kama basin are in good agreement with the idea of stabilization systems of gene pools [Artamonova, Makhrov, 2007 (Artamovova, Makhrov, 2007)]. When colonizing new reservoirs, at initial stages, processes of destabilization of the genetic structure are observed (a significant effect of selection, expressed in significant deviations of allele frequencies of the majority of genetic loci). For kilka, this process was traced in the early 2000’s with the settlement in the Rybinsk reservoir. Later, according to our observations, the genetic indicators stabilize (allele frequencies approach equilibrium ones), local stocks (intrapopulation groups) begin to form; kilka becomes an organic component of the ecosystem of the reservoir. Maintenance of the stable state of the population is provided by balancing selection, which retains a rather high level of polymorphism in the Upper Volga populations of common kilka [Karabanov, 2013b]. In the northern reservoirs, significantly differing in all respects from the waterbodies of the historical part of kilka’s range, regular changes occur in fish existence conditions, primarily associated with wintering. Accordingly, the adaptive value of different genotypes is constantly changing. The fairly high heterozygosity of this short-cycle species is supported by the presence of balancing selection during some seasons, which is replaced by disruptive selection, providing larger genotypic diversity in the population, which, in turn, can be changed by directional selection for a particular genotype. Thus, the population maintains a state of high fitness for living conditions, and a broad reaction rate is provided due to less adapted to the given conditions, but potentially adaptive genotypes - carriers of possible modifications that allow the organism to survive when the environmental factors outside the zone of the physiological optimum change [Shishkin, 1987]. On the whole, the consideration of adaptation transformations in the case of biological invasions makes it possible to study this process “here and now” in natural conditions, thanks to which (along with the involvement of theoretical constructions based on laboratory studies), a better understanding of microevolutionary transformations in marginal populations of animals is achieved.

> *Authors are deeply grateful to PhD A.A. Makhrov (IEEP RAS), prof. M.V. Mina (IDB RAS) and prof. RAS A.A. Kotov (IEEP RAS) for valuable advices on the peculiarities of modern taxonomy and phylogeography of animals.*
>
> *This research was performed in the framework of the state assignment of FASO Russia (theme No. AAAA-A18-118012690102-9) (sampling), supported in part by RFBR (project No. 17-05-00782-a) (genetic studies).*

## REFERENCES

April J., Mayden R.L., Hanner R.H., Bernatchez L. Genetic calibration of species diversity among North America’s freshwater fishes // Proceedings of the National Academy of Sciences. 2011. V. 108. No. 26. P. 10602–10607.

Artamonova V.S., Makhrov A.A. Geneticheskie sistemy kak reguliatory protcessov adaptatcii i vidoobrazovaniia (k sistemnoi teorii mikroevoliutcii) [Genetic systems as regulators of processes and speciation (to the system theory of microevolution)] / Sovremennye problemy biologicheskoi evoliutcii. M.: Gosudarstvennyi Darvinovskii muzei, 2007. S. 381–403. [In Russian]

Artamonova V.S., Makhrov A.A., Karabanov D.P., Rolskiy A.Y. et al. Hybridization of beaked redfish (Sebastes mentella) with small redfish (Sebastes viviparus) and diversification of redfish (Actinopterygii: Scorpaeniformes) in the Irminger Sea // Journal of Natural History. 2013. V. 47. №25–28. P. 1791–1801.

Askeyev O.V., Askeyev I.V., Ananin A.N., Tishin D.V. Obnaruzhenie deviatiigloi koliushki (Pungitius pungitius Linnaeus, 1758) v basseine r. Kamy (g. Nizhnekamsk, Respublika Tatarstan) [A record of nine-spined stickleback (Pungitius pungitius Linnaeus, 1758) in the Kama river basin (Nizhnekamsk, Republic Tatarstan)] // Povolzhskii ekologicheskii zhurnal. 2010. № 1. S. 103–106. [In Russian]

Atlas presnovodnykh ryb Rossii [Atlas of Freshwater Fishes of Russia]. Ed.: Yu.S. Reshetnikov. Moscow: Nauka, 2003a. V. 1. – 379 s. [In Russian]

Atlas presnovodnykh ryb Rossii [Atlas of Freshwater Fishes of Russia]. Ed.: Yu.S. Reshetnikov. Moscow: Nauka, 2003b. V. 1. – 253 s. [In Russian]

Bekker E.I., Karabanov D.P., Galimov Y.R., Kotov A.A. DNA barcoding reveals high cryptic diversity in the North Eurasian Moina species (Crustacea: Cladocera) // PLoS ONE. 2016. V. 11. No. 8. P. e0161737.

Bekker E.I., Karabanov D.P., Galimov Y.R., Haag C.R., Neretina T.V., Kotov A.A. Phylogeography of Daphnia magna Straus (Crustacea: Cladocera) in Northern Eurasia: Evidence for a deep longitudinal split between mitochondrial lineages //PLoS ONE. 2018. Vol. 13. No. 3. P. e0194045.

Biological invasions in changing ecosystems: vectors, ecological impacts, management and predictions. Ed.: J. Canning-Clode. Germany, Berlin: De Gruyter, 2015. – 488 p.

Biologicheskie invazii v vodnykh i nazemnykh ekosistemakh [Biological invasions in aquatic and terrestrial ecosystems]. Eds.: A.F. Alimov, N.G. Bogutckaia. M.-SPb.: KMK – ZIN RAN, 2004. – 436 s.

Boldyrev V.S., Bogutskaya N.G. Revision of the tadpole-gobie of the genus Benthophilus (Teleostei: Gobiidae) // Ichthyological Exploration of Freshwaters. 2007. V. 18. No. 1. P. 31–96.

Borovikova E.A. Filogeografiia riapushek Coregonus albula (L.) i C. sardinella (Valenciennes) Evropei`skogo Severa Rossii [Phylogeography of the vendace species Coregonus albula (L.) and C. sardinella (Valenciennes) of the European North of Russia]. Avtoref. dis. … kand. biol. nauk. M., 2009. – 24 s. [In Russian]

Borovikova E.A., Alekseeva Y.I., Schreider M.J., et al. Morphology and genetics of the ciscoes (Actinopterygii: Salmoniformes: Salmonidae: Coregoninae: Coregonus) from the Solovetsky Archipelago (White Sea) as a key to determination of the taxonomic position of ciscoes in northeastern Europe // Acta Ichthyologica et Piscatoria. 2013. V. 43. No. 3. P. 183–194.

Borovikova E.A., Makhrov A.A. Sistematicheskoe polozhenie i proishozhdenie sigov (Coregonus) Evropy: morfoekologicheskii podkhod [Taxonomy and origin of whitefish and ciscoes (Coregonus) in Europe: a morphoecological approach] // Trudy Karelskogo nauchnogo centra RAN. 2013. No. 6. S. 105–115.

Ekologicheskie problemy Verkhnei Volgi [Ecological problems of the Upper Volga]. Ed.: A.I. Kopylov. Yaroslavl: Izdatelstva YaGTU, 2001. – 427 s. [In Russian]

Eschmeyer W.N., Fricke R., van der Laan R. Catalog of fishes: genera, species, references. (http://researcharchive.calacademy.org/research/ichthyology/catalog/fishcatmain.asp). Electronic version accessed in Oct-2017.

Evlanov I.A., Kozlovskii S.I., Antonov P.I., Kadastr ryb Samarskoi oblasti [The inventory of fishes of the Samara region], Tolyatti: IEVB RAN, 1998. – 222 s. [In Russian]

Froese R., Pauly D., Editors. FishBase. World Wide Web electronic publication. (www.fishbase.org). Electronic version accessed in 10/2017.

Freyhof J. Diversity and Distribution of Freshwater Gobies from the Mediterranean, the Black and Caspian Seas / The Biology of Gobies. Ed.: B.G. Kapoor. USA, Boca Raton: CRC Press, 2011. P. 279–288

Galanin I.F. On expansion of goby fishes (Neogobius and Proterorhinus) in shallow shore areas of Kuybyshev Reservoir, Russia // Russian Journal of Biological Invasions. 2012. V. 3. No. 2. P. 101–104.

Gavlena F.K. Bychok-golovach Neogobius kessleri (Gunther) v Volgogradskom vodokhranilishche [Bighead goby Neogobius kessleri (Gunther) in the Volgograd reservoir] // Voprosy ikhtiologii. 1977. T. 17. No. 2. S. 359–360. [In Russian]

Gavlena F.K. Kaspiiskii bychok-krugliak Neogobius melanostomus affinis (Eichwald) – novyi element ikhtiofauny Srednei Volgi [Round goby Neogobius melanostomus affinis (Eichwald) – a new element of the ichthyofauna of the Middle Volga] // Biologiia vnutrennikh vod (informatcionnyi biulleten). 1970. No. 6. S. 44–45. [In Russian]

Gavlena F.K. Zvyozdchataia pugolovka Benthophilus stellatus (Sauvage) v Kuibyshevskom vodokhranilishche [Stellate tadpole-goby Benthophilus stellatus (Sauvage) in the Kuibyshev Reservoir] // Voprosy ikhtiologii. 1973. T. 13. No. 1. S. 174–175. [In Russian]

de Gelas K., de Meester L. Phylogeography of Daphnia magna in Europe // Molecular ecology. 2005. V. 14. No. 3. P. 753–764.

Gerasimov Yu.V., Smirnov A.K., Kodukhova Y.V. Assessment of possible causes of changes in abundance and sexual structure in populations of Prussian Carp (Carassius auratus gibelio Bloch, 1783) // Inland Water Biology. 2018. V. 11. № 1. P. 70–79.

Golovanov V.K., Kapshai D.S., Gerasimov Yu.V., et al. Thermopreference and thermostabilty of the Amur sleeper juveniles Perccottus glenii in autumn // Journal of Ichthyology. 2013. V. 53. No. 3. P.240–244.

Hellmuth M. Biologically feasible gene trees, reconciliation maps and informative triples // Algorithms for Molecular Biology. 2017. V. 12. N23.

Hubert N., Meyer C.P., Bruggemann H.J., et al. Cryptic diversity in Indo-Pacific coral-reef fishes revealed by DNA-barcoding provides new support to the Centre-of-Overlap hypothesis // PLoS ONE. 2012. V. 7. No. 3. P. e28987.

Opredelitel ryb i bespozvonochnykh Kaspiiskogo moria. T. 1. Ryby i molliuski. [Identification keys for fish and invertebrates. Volume 1. Fish and molluscs.] Ed.: N.V. Aladin. SPb. – M.: KMK, 2013. – 543 p

Karabanov D.P. Biohimicheskii polimorfizm v populiatciiakh chernomorsko-kaspiiskoi tiulki Clupeonella cultriventris (Nordmann, 1840) iz raznykh chastei areala [Biochemical polymorphism in populations of the common kilka Clupeonella cultriventris (Nordmann, 1840) from different parts of the area] // Vestnik AGTU. Serija: Rybnoe hozjajstvo. 2013b. No. 2. S. 159–165. [In Russian]

Karabanov D.P. Geneticheskie adaptatcii chernomorsko-kaspiiskoi tiulki Clupeonella cultriventris (Nordmann, 1840) (Actinopterygii: Clupeidae) [Genetical adaptation of common kilka Clupeonella cultriventris (Nordmann, 1840) (Actinopterygii: Clupeidae)]. Voronezh: Izdatelstvo “Nauchnaia kniga”, 2013a. – 179 s. [In Russian]

Karabanov D.P., Bazarov M.I., Kodukhova Yu.V. First record of Round Goby Neogobius melanostomus (Perciformes, Gobiidae) in the Uglich Reservoir (Volga River basin) // Inland Water Biology. 2014. V. 7. No. 4. P. 406–407.

Kasyanov A.N., Goroshkova T.V. Morphological features of the Amur sleeper (Perccottus glenii, Perciformes, Eleotridae) introduced into water bodies of European Russia // Contemporary Problems of Ecology. 2012. V. 5. No. 1. P. 58–70.

Kasyanov A.N., Klevakin A.A. Stellate tadpole-goby Benthophilus stellatus (Sauvage, 1874) in the Cheboksary Reservoir // Russian Journal of Biological Invasions. 2011. V. 2. No. 4. P. 242–244.

Kiryukhina N.A. Molecular and genetic variability in populations of Syngnathus nigrolineatus Eichwald 1831 and ways of expansion in the Volga River basins on the basis of mitochondrial DNA sequence analysis // Russian Journal of Biological Invasions. 2013. V. 4. No. 4. P. 249–254.

Kiyashko V.I., Karabanov D.P., Yakovlev V.N., Slynko Yu.V. Formation and development of the Black Sea-Caspian kilka Clupeonella cultriventris (Clupeidae) in the Rybinsk reservoir // Journal of Ichthyology. 2012. Vol.52. No.8. P. 537–546.

Klevakin, A.A., Blinov, Yu.V., Minin, A.E., et al. Rybolovstvo v Nizhegorodskoi oblasti [Fishing in the Nizhny Novgorod region], Nizhny Novgorod: Cheboksarskaya tipografiya No. 1. 2005. – 96 s. [In Russian]

Koblitckaya A.F. Opredelitel molodi presnovodnykh ryb [Handbook of juvenile freshwater fish]. M.: Lyogkaia i pishchevaia promyshlennost, 1981. – 208 s. [In Russian]

Kodukhova Y.V. Yearly variations of impact of natural hybrids of bream and roach (Abramis brama (L.) x Rutilus rutilus (L.)) in Rybinsk Reservoir // Russian Journal of Biological Invasions. 2011. V. 2. No. 2-3. P. 204–208.

Kodukhova Yu.V., Borovikova E.A., Karabanov D.P. First record of stellate tadpole goby Benthophilus stellatus (Sauvage, 1874) (Actinopterygii: Gobiidae) in the Rybinsk Reservoir // Inland Water Biology. 2016. V. 9. No. 4. P. 428–430.

Kotov A.A., Karabanov D.P., Bekker E.I., et al. Phylogeography of the Chydorus sphaericus group (Cladocera: Chydoridae) in the Northern Palearctic // PLoS ONE. 2016. V. 11. No. 12. P. e0168711.

Kottelat M., Freyhof J. Handbook of European freshwater fishes. Cornol: Publications Kottelat, 2007. 646 p.

Kozlovskaya S.I. Bychki v Sarahtovskom vodokhranilishche [Gobiid fishes in Saratov reservoir] // Voprosy ikhtiologii. 1997. T. 37. No. 3. S. 420. [In Russian]

Levin B.A., Simonov E.P., Ermakov O.A., Levina M.A. et al. Phylogeny and phylogeography of the roaches, genus Rutilus (Cyprinidae), at the Eastern part of its range as inferred from mtDNA analysis // Hydrobiologia. 2017. V. 788. No. 1. P. 33–46.

Martemyanov V.I., Borisovskaya E.A. Indices of hydromineral metabolism in tyulka (Clupeonella cultriventris; Clupeiformes, Clupeidae) introduced in the Rybinsk Reservoir in comparison to aboriginal and marine fish species // Russian Journal of Biological Invasions. 2010. V. 1. No. 3. P. 187–193.

Mina M.V., Reshetnikov Y.S., Dgebuadze Y.Y. Taxonomic novelties and problems for users // Journal of Ichthyology. 2006. V. 46. No. 4. P. 476–480.

Moskalkova K.I. Ekologicheskie i morfo-fiziologicheskie predposylki k rasshireniiu areala u bychka-krugliaka Neogobius melanostomus v usloviiakh antropongennogo zagriazneniia vodoemov [Ecological and morpho-physiological prerequisites for expanding the range of the round goby Neogobius melanostomus in conditions of anthropogenic pollution of water bodies] // Voprosy ikhtiologii. 1996. T. 36. No. 5. S. 615–621. [In Russian]

Nei M., Kumar S. Molecular evolution and phylogenetics. NY: Oxford University Press, 2000. – 333 p.

Neilson M.E., Stepien C.A. Escape from the Ponto-Caspian: Evolution and biogeography of an endemic goby species flock (Benthophilinae: Gobiidae: Teleostei) // Molecular Phylogenetics and Evolution. 2009. V. 52. No. 1. P. 84–102.

Nelson, J.S., Grande T.C., Wilson M.V.H. Fishes of the World. Fifth edition. Hoboken, New Jersey, USA: John Wiley and Sons, 2016. – 707 p.

Prentis P.J., Wilson J.R.U., Dormontt E.E., et al. Adaptive evolution in invasive species // Trends in Plant Science. 2008. V. 13. No. 6. P. 288–294.

Pravdin I.F. Rukovodstvo po izucheniyu ryb (preimushchestvenno presnovodnykh) [Fish Study Manual (Predominantly Freshwater Species)], Moscow: Pishchevaya Promyshlennost, 1966. – 376 s. [In Russian]

Reshetnikov A.N., Ficetola G.F. Potential range of the invasive fish rotan (Perccottus glenii) in the Holarctic // Biological Invasions. 2014. V. 13. No. 12. P. 2967–2980.

Reshetnikov Yu.S., Kotlyar A.N., Rass T.S., Shatunovskii M.I. Piatiiazychnyi slovar nazvanii zhivotnykh. Ryby. Latinskii, russkii, angliiskii, nemetckii, frantcuzskii [Five-language dictionary of animal names. Fish. Latin, Russian, English, German, French]. M.: Russkii iazyik, 1989. – 355 s. [In Russian]

Ryby Rybinskogo vodokhranilishcha: populiatcionnaia dinamika i ekologiia [Fish of Rybinsk Reservoir: population dynamics and ecology]. Ed.: Yu.V. Gerasimov. Yaroslavl: Filigran, 2015. – 418 s. [In Russian]

Ryby v zapovednikakh Rossii [Fish in the reserves of Russia]. Ed.: Yu.S. Reshetnikov. Moskow: KMK, 2010. T. 1. – 627 s. [In Russian]

Semyonov D.Yu. Dannye o morfologii i biologii goloveshki-rotana Perccottus glenii Dybowski, 1877 (Perciformes, Eleotrididae) Kuibyshevskogo vodokhranilishcha [Data of the morphology and biology of rotan Perccottus glenii Dybowski, 1877 (Perciformes, Eleotrididae) of the Kuibyshev reservoir] // Yug Rossii: ekologiia, razvitie. 2010. № 3. P. 88–93. [In Russian]

Shakirova F.M., Severov Y.A., Latypova V.Z. Modern composition of alien fish species in the Kuybyshev reservoir and possible introduction of new representatives into its ecosystem // Russian Journal of Biological Invasions. 2015. V. 6. No. 4. P. 278–291.

Skomorokhov M.O. Caspian bighead goby Neogobius gorlap Iljin in Berg, 1949 (Gobiidae, Pisces) – a new invader species in the Moscow River // Russian Journal of Biological Invasions. 2016. V. 7. No. 3. P. 297–301.

Slynko Y.V., Borovikova E.A., Gurovskii A.N. Phylogeography and origin of freshwater populations of tubenose gobies of genus Proterorhinus (Gobiidae: Pisces) in Ponto-Caspian Basin // Russian Journal of Genetics. 2013. V. 49. No. 11. P. 1144–1154.

Slynko Y.V., Karabanov D.P., Stolbunova V.V. Genetic analysis of the intraspecific structure of kilka Clupeonella cultiventris (Nordmann, 1840) (Actinopterigii: Clupeidae) // Doklady Biological Sciences. 2010. V. 433. No. 1. P. 261–263.

Slynko Yu. V., Tereshchenko V. G. Ryby presnyh vod ponto-kaspijskogo bassejna raznoobrazie faunogenez dinamika populyacij mekhanizmy adaptacij [Freshwater fish of the Ponto-Caspian basin (Variety, faunogenesis, population dynamics, adaptation mechanisms)]. M: Izdatelstvo “Poligraf-Plus”, 2014. – 328 p.

Sokolov L.I., Tcepkin E.A. Istoricheskii obzor antropogennykh izmenenii ikhtiofauny rek Centralnogo regiona Rossii (na primere basseina Moskvy-reki i drugikh rek Podmoskovia) [Historical review of anthropogenic changes in the ichthyofauna of rivers in the Central region of Russia (on the example of the Moscow River basin and other rivers of the Moscow region)] // Voprosy ikhtiologii. 2000. T. 40. No. 2. S. 166–175. [In Russian]

Sorokin P.A., Medvedev D.A., Vasil’ev V.P., Vasil’eva E.D. Further studies of mitochondrial genome variability in Ponto-Caspian Proterorhinus species (Actinopterygii: Perciformes: Gobiidae) and their taxonomic implications // Acta Ichthyologica et Piscatoria. 2011. V. 41. No. 2. P. 95–104.

Stolbunov I.A., Malin M.I., Karabanov D.P. Finding round goby Neogobius melanostomus (Pallas, 1814) in the Rybinsk Reservoir // Inland Water Biology. 2013. V. 6. No. 4. P. 365–367.

Svetovidov A.N. Clupeidae / Check-list of the Fishes of the North-eastern Atlantic and of the Mediterranean J.-C. Hureau, T. Monod (eds.). V. 1. Paris: UNESCO, 1973. P. 99–109.

Tereshchenko V.G., Tereshchenko L.I. Potentcialnaia skorost rosta chislennosti populiatcii snetka Osmerus eperlanus (L.) v usloviiakh Kuibyshevskogo vodokhranilishcha [Specific growth rate of smelt Osmerus eperlanus (L.) population in Kuibyshev reservoir] // Transactions of IBIW RAS. 2017. No. 80(83). P. 86–94. [In Russian]

Tciplakov E.P. Rasshirenie arealov nekotorykh vidov ryb v sviazi s gidrostroitelstvom na Volge i akclimatizatcionnymi rabotami [Expansion of ranges of certain fish species due to the hydraulic works on the Volga River and acclimatization activity] // Voprosy ikhtiologii. 1974. T. 14. No. 3. S. 396–405.

Swenson K.M., El-Mabrouk N. Gene trees and species trees: irreconcilable differences // BMC Bioinformatics. V. 13. P. S15.

Ward R.D., Hanner R., Hebert P.D. The campaign to DNA barcode all fishes, FISH-BOL // Journal of Fish Biology. 2009. V. 74. No. 2. P. 329–356.

Zworykin D.D., Pashkov A.N. Eight-striped cichlasoma – an allochthonous species of cichlid fish (Teleostei: Cichlidae) from Staraya Kuban Lake // Russian Journal of Biological Invasions. 2010. V. 1. No. 1. P. 1–6.

